# Natural variation in stochastic photoreceptor specification and color preference in *Drosophila*

**DOI:** 10.1101/153445

**Authors:** Caitlin Anderson, India Reiss, Cyrus Zhou, Annie Cho, Haziq Siddiqi, Ben Mormann, Cameron M. Avelis, Alan Bergland, Elijah Roberts, James Taylor, Daniel Vasiliauskas, Robert J. Johnston

## Abstract

Each individual perceives the world in a unique way, but little is known about the genetic basis of variation in sensory perception. Here we investigated natural variation in the development and function of the color vision system of *Drosophila*. In the fly eye, the random mosaic of color-detecting R7 photoreceptor subtypes is determined by stochastic expression of the transcription factor Spineless (Ss). Individual R7s randomly choose between Ss^ON^ or Ss^OFF^ fates at a ratio of 65:35, resulting in unique patterns but consistent proportions of cell types across genetically identical retinas. In a genome wide association study, we identified a naturally occurring insertion in a regulatory DNA element in the *ss* gene that lowers the ratio of Ss^ON^ to Ss^OFF^ cells. This change in photoreceptor fates shifts the innate color preference of flies from green to blue. The genetic variant increases the binding affinity for Klumpfuss (Klu), a zinc finger transcriptional repressor that regulates *ss* expression. Klu is expressed at intermediate levels to determine the normal ratio of Ss^ON^ to Ss^OFF^ cells. Thus, binding site affinity and transcription factor levels are finely tuned to regulate stochastic on/off gene expression, setting the ratio of alternative cell fates and ultimately determining color preference.

---

Organisms require a diverse repertoire of sensory receptor neurons to perceive a range of stimuli in their environments. Differentiation of sensory neurons often requires stochastic mechanisms whereby individual neurons randomly choose between different fates. Stochastic fate specification diversifies sensory neuron subtypes in a wide array of species including worms, flies, mice, and humans (Ressler et al. 1993; Roorda and Williams 1999; Troemel et al. 1999; Hofer et al. 2005; Johnston and Desplan 2010; Magklara and Lomvardas 2013; Alqadah et al. 2016; Viets et al. 2016). How naturally occurring changes in the genome affect stochastic mechanisms to alter sensory system development and perception is poorly understood. To address this question, we investigated natural variation in stochastic color photoreceptor specification in the *Drosophila* retina.

The fly eye, like the human eye, contains a random mosaic of photoreceptors defined by expression of light-detecting Rhodopsin proteins (Montell et al. 1987; Bell et al. 2007; Johnston and Desplan 2010; Viets et al. 2016). In flies, the stochastic on/off expression of Spineless (Ss), a PAS-bHLH transcription factor, determines R7 photoreceptor subtypes. Ss expression in a random subset of R7s induces ‘yellow’ (**y**R7) fate and expression of Rhodopsin4 (Rh4), whereas the absence of Ss in the complementary subset of R7s allows for ‘pale’ (**p**R7) fate and Rhodopsin3 (Rh3) expression (Fig. 1A)(Wernet et al. 2006; Johnston et al. 2011; Thanawala et al. 2013; Johnston and Desplan 2014). The on/off state of Ss in a given R7 also indirectly determines the subtype fate of the neighboring R8 photoreceptor. pR7s lacking Ss signal to pR8s to activate expression of blue-detecting Rhodopsin 5 (Rh5). **y**R7s expressing Ss do not send this signal, resulting in expression of green-detecting Rhodopsin 6 (Rh6) in yR8s (Fig 1A)(Franceschini et al. 1981; Montell et al. 1987; Zuker et al. 1987; Chou et al. 1996; Huber et al. 1997; Chou et al. 1999; Mikeladze-Dvali et al. 2005; Jukam and Desplan 2011; Hsiao et al. 2013; Johnston 2013; Jukam et al. 2013; Jukam et al. 2016).

**Figure 1.**
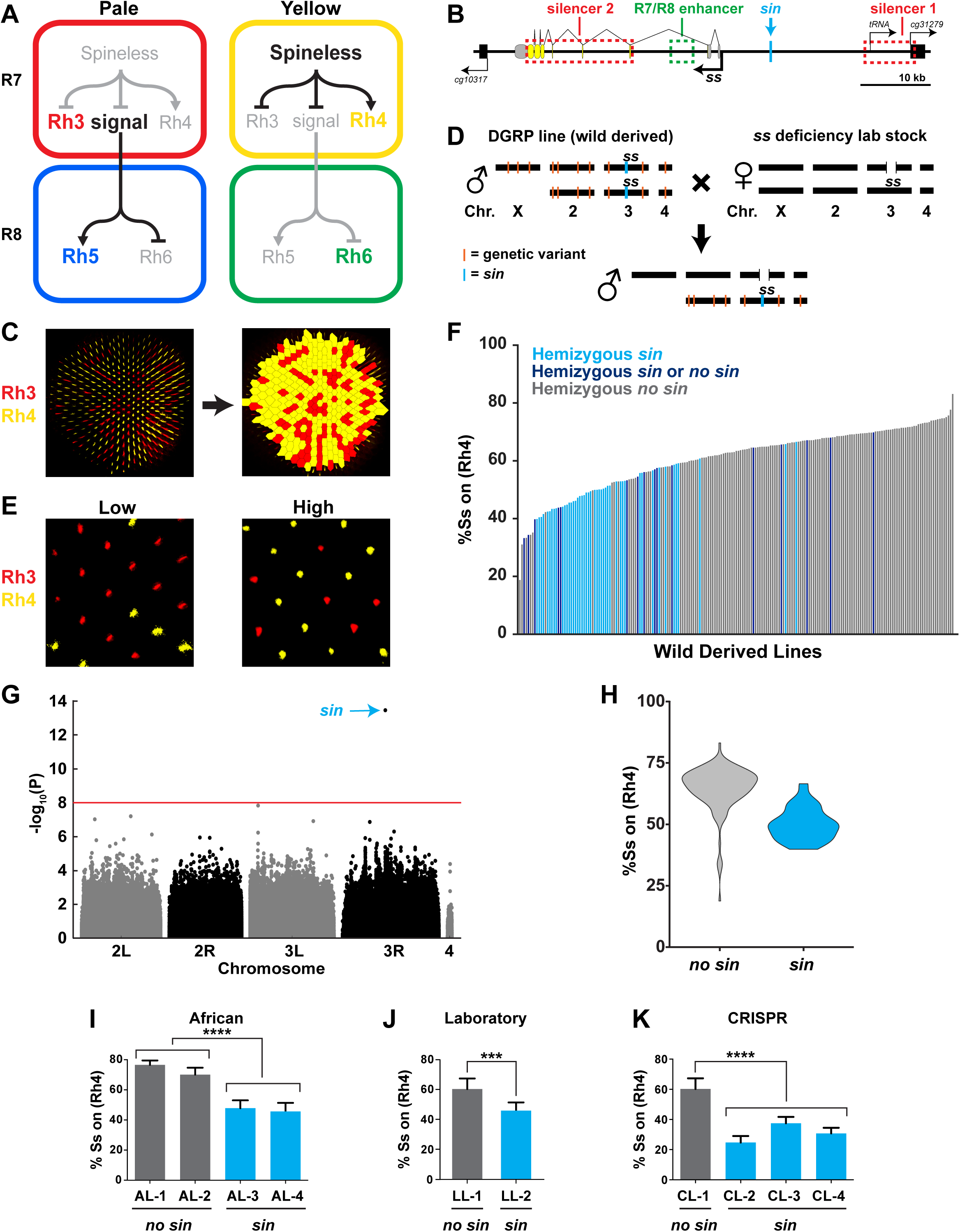
A naturally-occurring single base insertion in the *ss* locus (*sin*) lowers the ratio of Ss^ON^ to Ss^OFF^ R7s. A. R7 and R8 subtypes are determined by the on/off expression of Spineless (Ss). (Left) The absence of Ss allows Rh3 expression in pale R7s and Rh5 expression in pale R8s. (Right) Expression of Ss induces Rh4 expression in yellow R7s and Rh6 expression in yellow R8s. The signal by which Spineless mediates Rh5 vs. Rh6 expression in R8s is currently unknown. B. Schematic of the *ss* locus. Green dashed rectangle indicates *R7/R8 enhancer*; red dashed rectangles indicate *silencer 1* and *silencer 2*; blue line indicates Klu binding site; blue arrow indicates *<u>s</u>s <u>in</u>sertion/sin;* gray ovals represent untranslated exons; yellow ovals represent translated exons; black boxes indicate neighboring genes; arrows indicate transcriptional starts. C. Image of a whole mount fly retina. (Left) Stochastic distribution of R7s expressing Rh3 (Ss^OFF^) or Rh4 (Ss^ON^). (Right) An automated counting system identified and counted Rh3- and Rh4- expressing R7s. D. Crossing scheme: Wild-derived DGRP flies were crossed with *ss* deficiency flies, yielding progeny that were hemizygous at the *ss* locus. Orange lines indicate genetic variants; blue line indicates *sin* in *ss.* E. Representative images from progeny in (D) with low (left; DGRP-397) and high (right; DGRP-229) proportions of Ss^ON^ (Rh4) R7s. F. Ss^ON^ proportion varied across DGRP fly lines. *sin* was enriched in lines with a low proportion of Ss^ON^ R7s. Each bar represents a single DGRP line, and bars are arranged in rank order. Light blue bars indicate hemizygous *sin*. Dark blue bars indicate hemizygous *sin* or hemizygous *no sin* (original DGRP line was heterozygous *sin/no sin*). Gray bars indicate hemizygous *no sin*. G. GWAS identified *sin* as a genetic variant associated with Ss expression. Manhattan plot of the genetic variant p-values. Genetic variants above the red line (Bonferroni correction) are considered significant. Arrow indicates *sin*. H. *sin* was enriched in lines with a low proportion of Ss^ON^ R7s. Violin plot of DGRP lines with and without *sin*. I-K. Flies with *sin* displayed a lower proportion of Ss^ON^ R7s compared to flies without *sin.* AL indicates African lines; LL indicates laboratory lines; CL indicates lines in which *sin* was inserted with CRISPR. ^****^ indicates p<0.0001; ^***^ indicates p<0.001. Error bars indicate standard deviation (SD).

The stochastic decision to express Ss is made cell autonomously at the level of the *ss* gene locus via a random repression mechanism. The *R7/R8 enhancer* induces *ss* expression in all R7s, whereas two silencer regions (*silencer 1* and *2*) repress expression in a random subset of R7s (Fig. 1B)(Johnston and Desplan 2014).

Though the stochastic expression of Ss is binary (i.e. on or off) in individual R7s, it does not result in a simple 50:50 on/off ratio across the population of R7s in a given retina. In most isogenic lab stock flies, Ss is on in ~65% of R7s and off in ~35% (Fig. 1C)(Wernet et al. 2006; Johnston and Desplan 2014). Here, we find that the proportion of Ss^ON^ to Ss^OFF^ R7s varies greatly among fly lines derived from the wild. We performed a genome-wide association study (GWAS) and identified a single base pair insertion that increases the affinity of a DNA binding site for a transcriptional repressor, significantly reducing the Ss^ON^/Ss^OFF^ ratio. This genetic variant changes the proportion of photoreceptor subtypes and alters the innate color preference of flies.

## *sin* decreases the ratio of Ss^ON^ to Ss^OFF^ R7s

To determine the mechanism controlling the ratio of stochastic on/off Ss expression, we analyzed the variation in 203 naturally-derived lines collected from Raleigh, North Carolina (Drosophila Genetic Reference Panel (DGRP))(Mackay et al. 2012). We evaluated Rh4 and Rh3 expression as they faithfully report Ss expression in R7s (i.e. Ss^ON^ = Rh4; Ss^OFF^ = Rh3) (Fig. 1A)(Thanawala et al. 2013; Johnston and Desplan 2014). To facilitate scoring, we generated a semi-automated counting system to determine the Rh4:Rh3 ratio for each genotype (Fig. 1C).

To assess the variation in the DGRP lines attributable to the *ss* locus and limit the phenotypic contribution of recessive variants at other loci, we crossed each DGRP line to a line containing a ~200 kb deficiency covering the *ss* locus and analyzed Rh3 and Rh4 expression in the F1 male progeny (Fig. 1D). This genetic strategy generated flies hemizygous (i.e. single copy) for the wild-derived *ss* gene locus, heterozygous wild-derived/lab stock for the 2^nd^, 3^rd^, and 4^th^ chromosomes, and hemizygous lab stock for the X chromosome (Fig. 1D). While the lab stock expressed Ss (Rh4) in 62% of R7s under these conditions, expression among the DGRP lines varied significantly, ranging from 19 to 83% Ss^ON^ (Rh4) (Fig. 1E-F; **Supplemental Table 1**).

To identify the genetic basis of this variation, we performed a genome-wide association study (GWAS) using the Ss^ON^ (Rh4) phenotype data and inferred full genome sequences of the progeny of each DGRP line crossed with the *ss* deficiency line. We performed an association analysis and identified a single base pair insertion within the *ss* locus (“*<u>s</u>s* <u>in</u>sertion” or “*sin*”) that was significant (p<10^-13^) after Bonferroni correction (Fig. 1G). *sin* was enriched in DGRP lines with a low ratio of Ss^ON^ to Ss^OFF^ R7s (Fig. 1F and H).

We next confirmed the regulatory role of *sin.* Naturally derived lines from Africa that are homozygous for *sin* displayed a decrease in the proportion of Ss^ON^ (Rh4) R7s compared to lines from Africa lacking *sin* (Fig. 1I)(Lack et al. 2015). We identified *sin* on a balancer chromosome (*TM6b*) in a lab stock that similarly displayed a decrease in the proportion of Ss^ON^ (Rh4) R7s when *ss* was hemizygous (Fig. 1J). To definitively test the role of *sin*, we used CRISPR to insert *sin* into a lab stock. Flies hemizygous for CRISPR *sin* alleles displayed a significant decrease in the proportion of Ss^ON^ (Rh4) R7s (Fig. 1K). Thus, *sin* causes a decrease in the ratio of Ss^ON^ to Ss^OFF^ R7s.

## *Sin* shifts innate color preference from green to blue

As *sin* alters the proportion of color-detecting photoreceptors, we hypothesized that it would also change color detection and preference. When presented with two light stimuli in a T-maze (Tully and Quinn 1985), flies will phototax toward the light source that they perceive as more intense (Fig. 2A)(McEwen 1918; Heisenberg and Wolf 1984; Choe and Clandinin 2005). The absorption spectra of Rh3 and Rh4 significantly overlap in the UV range (Feiler et al. 1992), complicating behavioral assessment of color preference caused by differences in R7 photoreceptor ratios. Instead, we focused on the perception of blue light by Rh5 and green light by Rh6 in the R8 photoreceptors, as these Rhodopsins have more distinct absorption spectra (Salcedo et al. 1999). Because R8 fate is coupled to R7 fate(Chou et al. 1996) (Fig. 1A), we predicted that flies with *sin* would have a low ratio of Rh6- to Rh5-expressing R8s and would consequently prefer blue light, while flies without *sin* would have a higher ratio of Rh6- to Rh5-expressing R8s and would instead prefer green light. Indeed, DGRP lines containing *sin* preferred blue light, while DGRP lines lacking *sin* preferred green light (Fig 2A-C; **Supplemental Table 2**), showing that *sin* changes innate color preference in flies.

**Figure 2.**
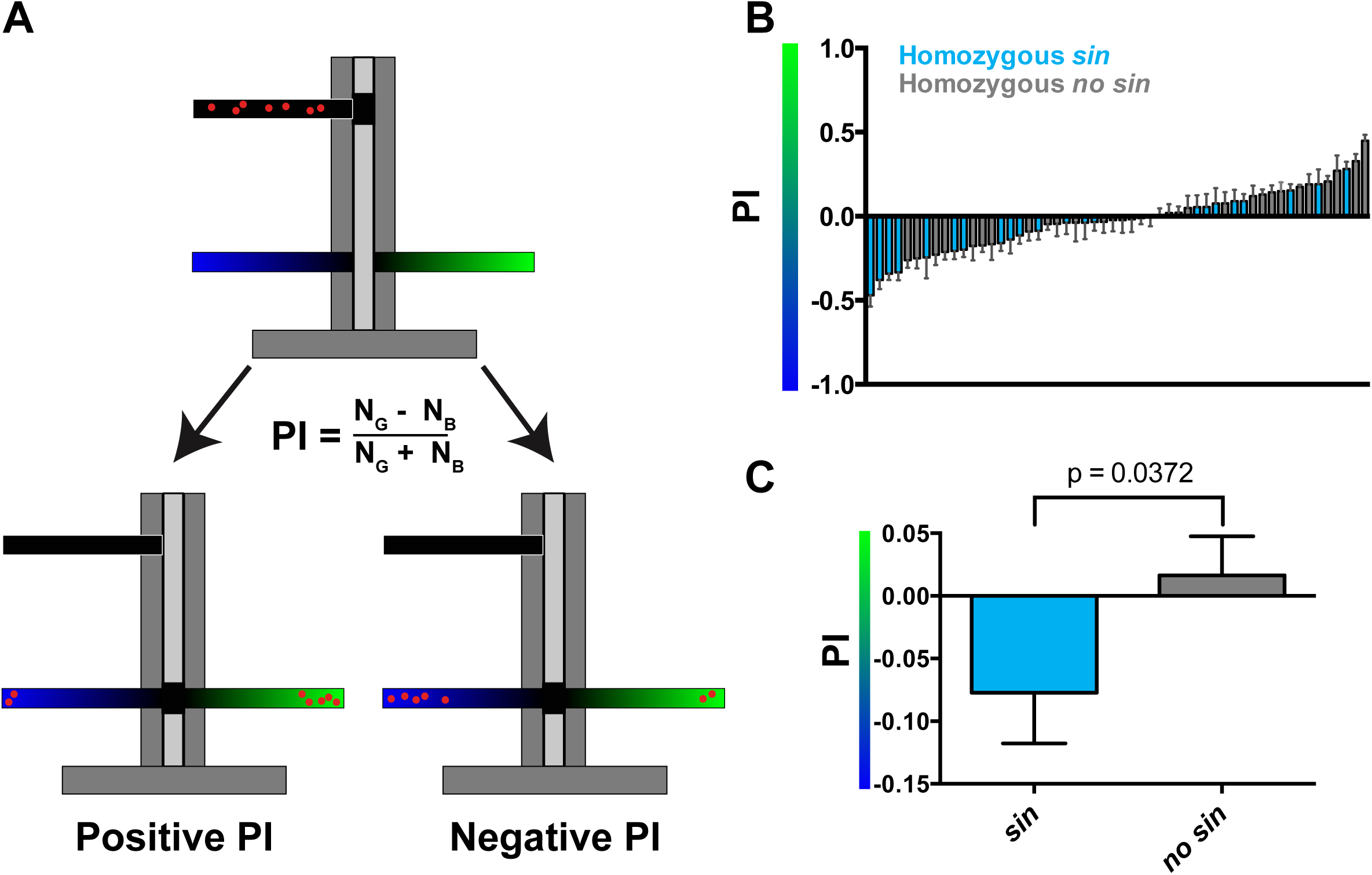
Wild-derived flies with *sin* display a shift in innate color preference from green to blue. A. Schematic of T-maze apparatus. Red dots represent flies. PI = Preference Index. Positive PI indicates preference for green light; negative PI indicates preference for blue light. NG: number of flies on green side; NB: number of flies on blue side. B-C. Flies with *sin* preferred blue light compared to flies without *sin* that preferred green light. Color of bar indicates genotype of DGRP line. Light blue bars indicate homozygous *sin*. Dark blue bars indicate heterozygous *sin* / *no sin.* Gray bars indicate homozygous *no sin*. Error bars indicate standard error of the mean (SEM). B. PIs for individual DGRP lines with and without *sin.* C. Averages of PIs for DGRP lines with and without *sin.*

## *sin* increases the binding affinity for the Klumpfuss transcription factor

*sin* is a single base pair insertion within a previously uncharacterized non-coding region of the *ss* locus located ~7 kb upstream of the transcriptional start (Fig. 1B and Fig. 3A). To identify *trans* factors whose binding might be affected by *sin*, we searched for binding motifs neighboring *sin* in bacterial one-hybrid (B1H)(Zhu et al. 2011; Enuameh et al. 2013) and SELEX-seq datasets(Nitta et al. 2015). *sin* lies in a predicted binding site for the zinc finger transcription factor Klumpfuss (Klu), the fly homolog of Wilms’ Tumor Suppressor Protein 1 (WT1) (Fig. 3B, **S1A**)(Klein and Campos-Ortega 1997; Yang et al. 1997). This region is perfectly conserved across 21 *Drosophila* species covering 50 million years of evolution, consistent with a critical regulatory role (Fig. 3C, **S1B-C**).

**Figure 3.**
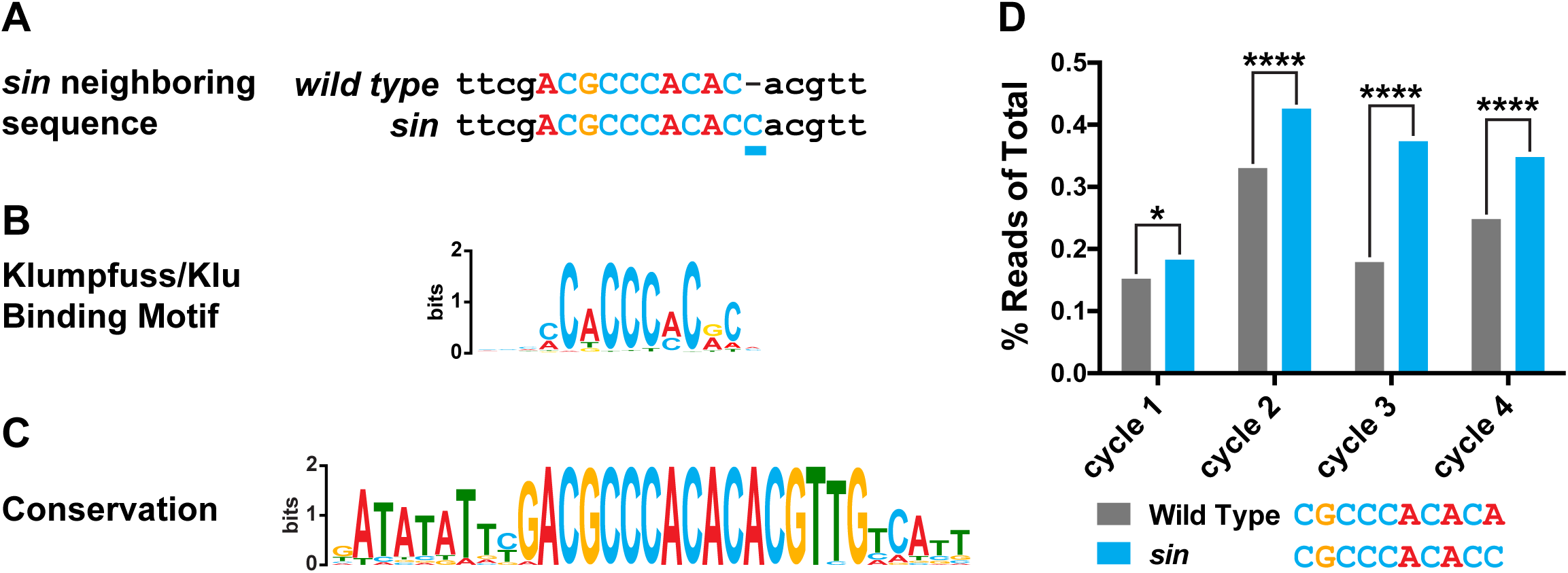
sin increases the binding affinity for the transcription factor Klumpfuss. A. sin is a single base pair insertion of a C at Chr. 3R: 16,410,775 (release 6). Underline indicates sin. B. sin affects a predicted binding site for the transcription factor Klumpfuss (Klu). Binding site predicted from B1H. C. The Klu site is perfectly conserved across 21 species of Drosophila covering 50 million years of evolution. Conservation logo of the Klu site and neighboring sequence in the ss locus. Height of bases indicates degree of conservation. D. sin increased Klu binding affinity in vitro. Quantification of the number of reads for the Klu site with and without sin in four cycles of SELEX-seq. ^*^ indicates p<0.05 in cycle 1; ^****^ indicates p<0.0001 in cycles 2-4.

Since *sin* does not affect the core Klu DNA binding motif (MCWCCCMCRC), we predicted that *sin* would not ablate binding, but would rather alter the affinity of Klu for the site. To evaluate the effect of *sin* on Klu binding, we analyzed available SELEX-seq binding data(Nitta et al. 2015). The number of reads containing the Klu binding site with *sin* (CGCCCACACC) was significantly higher than without *sin* (CGCCCACACA) (Fig. 3D), and thus, Klu binds sequences with *sin* better than those without it. Considering the frequency of 10-mers as a measure of site preference, we found that 506 10-mers (0.10%) have frequencies greater than the Klu site without *sin,* whereas only 366 10-mers (0.07%) have frequencies greater than the Klu site with *sin*. Together, *sin* increases the binding affinity of the Klu site.

## Klu lowers the Ss^ON^/Ss^OFF^ ratio in R7s

As *sin* increases Klu binding affinity and decreases Ss expression frequency, we predicted that Klu acts as a repressor of stochastic *ss* expression in R7s. Consistent with this hypothesis, Klu/WT1 has been shown in other systems to be a transcriptional repressor (Drummond et al. 1992; McDonald et al. 2003; Kaspar et al. 2008). Further, we found that Klu was expressed in all R7s in larval eye imaginal discs (Fig. 4A-B)(Wildonger et al. 2005). Since Klu is a repressor, we predicted that increasing Klu levels would cause a decrease in the proportion of Ss^ON^ R7s whereas decreasing or completely ablating Klu would cause an increase in the proportion of Ss^ON^ R7s. Indeed, increasing the levels of Klu in Klu-expressing cells (*klu>klu*), all photoreceptors (*eye>klu*), or specifically in all R7s (*R7>klu*) caused a decrease in the proportion of Ss^ON^ (Rh4) R7s (Fig. 4C-D). This decrease in the Ss^ON^/Ss^OFF^ ratio upon increasing Klu levels mimicked the effect of *sin*, consistent with *sin* increasing the binding affinity for the Klu repressor. Conversely, *klu* null and strong hypomorphic mutants displayed increases in the proportion of Ss^ON^ (Rh4) R7s (Fig. 4E-F). Since raising Klu levels decreased the Ss^ON^/Ss^OFF^ ratio and ablating *klu* increased the ratio, Klu is endogenously expressed at intermediate levels to determine the proportion of Ss-expressing R7s.

**Figure 4.**
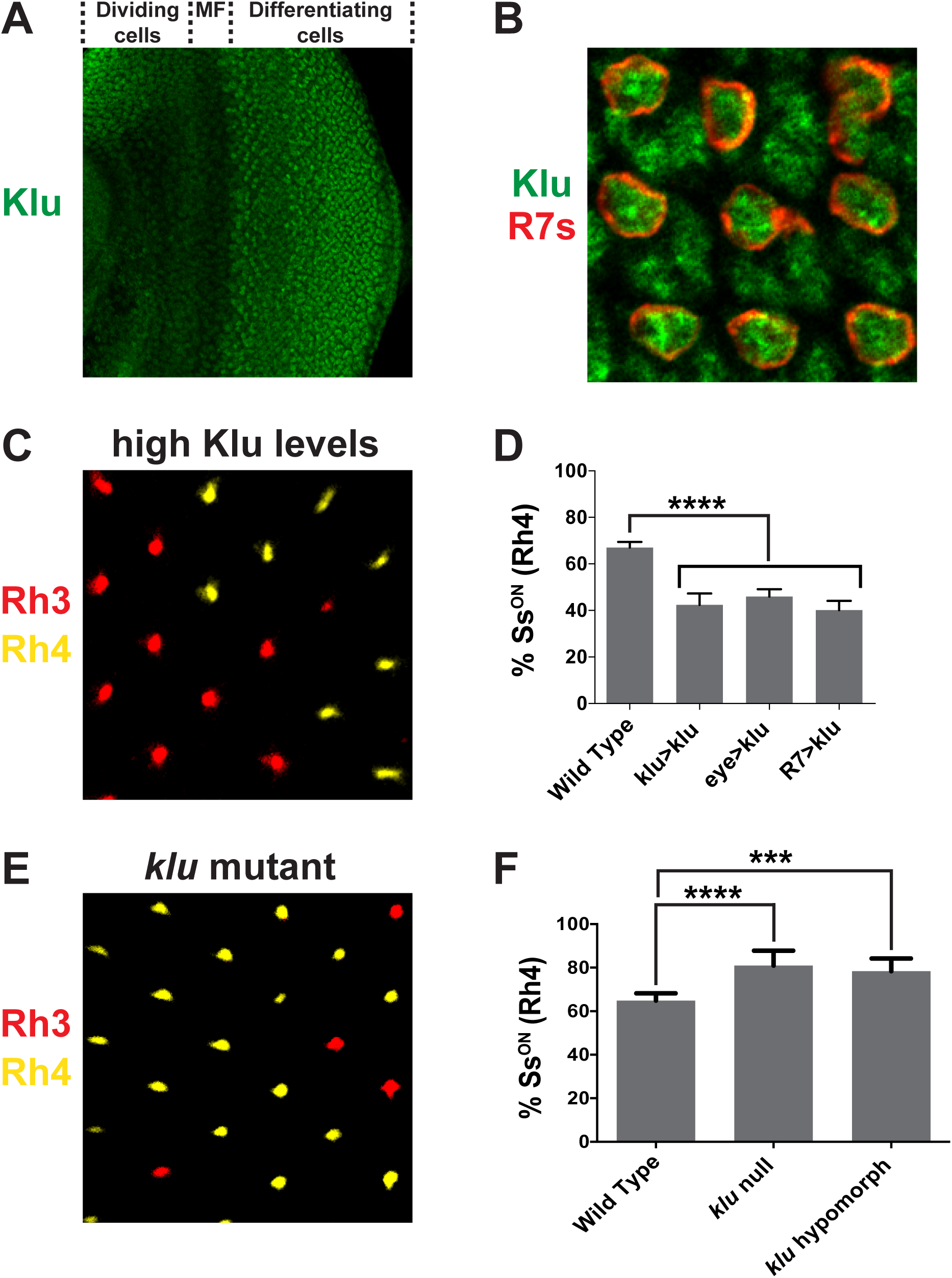
Levels of Klu determine the ratio of Ss^ON^/Ss^OFF^ R7s. A. Klu was expressed in the developing larval eye disc. MF indicates morphogenetic furrow. B. Klu was expressed in all R7s in the developing larval eye disc. Red indicates R7s marked by *pm181>Gal4, UAS>mcd8GFP.* C-D. Increasing Klu levels decreased the proportion of Ss^ON^ R7s. In C, representative image of R7s expressing Rh3 (Ss^OFF^) or Rh4 (Ss^ON^) in *eye>klu* flies. ^****^ indicates p<0.0001. Error bars indicate standard deviation (SD). E-F. *klu* loss of function mutants displayed increases in the proportion of Ss^ON^ R7s. In E, representative image of R7s expressing Rh3 (Ss^OFF^) or Rh4 (Ss^ON^) in *klu* hypomorphic flies. ^****^ indicates p<0.0001; ^***^ indicates p<0.001. Error bars indicate standard deviation (SD).

Our studies of wild-derived flies revealed significant variation in stochastic Ss expression. We identified *sin*, a single base pair insertion in the ~60 kb *ss* locus that dramatically lowers the Ss^ON^/Ss^OFF^ ratio by increasing the binding affinity of the transcriptional repressor Klu. This decrease in Ss expression frequency changes the proportions of color-detecting photoreceptors and alters innate color preference in flies.

*sin* appears to be a relatively new mutation in *D. melanogaster* populations. *sin* is absent among diverse drosophilid species spanning millions of years of divergence (**Fig. S1B-C**) and is segregating at an extremely low frequency among non-admixed African *D. melanogaster* lineages (**Fig. S2A-F**). *sin* likely rose to intermediate frequencies following *D. melanogaster’s* colonization of Europe about 10-15 thousand years ago (Li and Stephan 2006). *sin* continues to segregate at intermediate frequencies amongst North American populations (**Fig. S2A-F**), which were established within the last 150 years from mixtures of European and African populations (Bergland et al. 2016). The recent rise in frequency of *sin* suggests that it could be the target of natural selection, perhaps via modulation of innate color preference. We tested this model by assessing patterns of allele frequency differentiation among populations sampled world-wide and also through examination of haplotype homozygostiy surrounding *sin*. We compared these statistics at *sin* to the distribution of statistics calculated from several thousand randomly selected 1-2bp indel polymorphisms that segregate at ~25% in the DGRP. Curiously, *sin* did not deviate from genome-wide patterns (**Fig. S2G-J**) suggesting that it might be selectively neutral in contemporary *D. melanogaster* populations.

It is interesting that Rhodopsin expression varies so significantly in the wild, given the nearly invariant hexagonal lattice of ommatidia in the fly eye. Rhodopsins are G-protein coupled receptors (GPCRs), a class of proteins identified as a source of natural behavioral variation in worms, mice, and voles (Young et al. 1999; Yalcin et al. 2004; Bendesky et al. 2011). Dramatic differences in Rhodopsin expression patterns across insect species (Wernet et al. 2015) suggest that variation in the expression of GPCRs, rather than retinal morphology, may allow rapid evolution in response to environmental changes.

*sin* increases the binding affinity of a conserved Klu site, suggesting that the site is suboptimal or low affinity for Klu binding. Low affinity sites ensure the timing and specificity of gene expression (Jiang and Levine 1993; Gaudet and Mango 2002; Scardigli et al. 2003; Rowan et al. 2010; Ramos and Barolo 2013; Crocker et al. 2015; Farley et al. 2015; Crocker et al. 2016). Our studies have revealed a critical role for a low affinity binding site in the regulation of stochastic gene expression. The suboptimal Klu site, bound by endogenous levels of Klu, yields the normal 65:35 Ss^ON^/Ss^OFF^ ratio. Changing the affinity of the site or the levels of Klu alters the ratio of Ss^ON^/Ss^OFF^ cells. Together, we conclude that stochastic on/off gene expression is controlled by threshold levels of *trans* factors binding to low affinity sites.

Levels of Klu (analog input) determine the binary on/off ratio of Ss expression (digital output). In contrast, gene regulation is best understood in cases where levels of transcription factors (analog input) regulate the levels of target gene expression (analog output). The on/off nature of Ss expression suggests a cooperative mechanism whereby Klu acts with other factors to repress *ss.* The expression state of *ss* could be determined by the intrinsic variation in Klu levels. In this model, if Klu levels exceed a threshold, *ss* is off, and if Klu levels are below the threshold, *ss* is on. Alternatively, Klu levels could set the threshold for a different gene regulatory mechanism, such as DNA looping or heterochromatin spreading. The requirement of two silencers for repression is consistent with a role for DNA looping in stochastic *ss* expression (Johnston and Desplan 2014). Klu levels could shift the balance between DNA looping states that determine on or off *ss* expression.

Cell fate specification is commonly thought of as a reproducible process whereby cell types uniformly express specific batteries of genes. This reproducibility is often the result of high levels of transcription factors binding to high affinity sites, far exceeding a regulatory threshold, yielding expression of target genes in all cells of a given type. In contrast, the stochastic on/off expression of Ss requires finely tuned levels of regulators binding to low affinity sites. We predict that fine tuning of binding site affinities and transcription factor levels will emerge as a common mechanistic feature that determines the ratio of alternative fates in stochastic systems.

## Materials and Methods – (also see Supplemental Materials and Methods)

### *Drosophila* genotypes and stocks

Flies were raised on standard cornmeal-molasses-agar medium and grown at 25°C.

### Antibody Staining

Adult retinas and larval eye discs were dissected as described (Hsiao et al. 2012).

### Quantification of Expression

Frequency of Rh3 (Ss^OFF^) and Rh4 (Ss^ON^) expression in R7s was scored in adults. Six or more retinas were scored for each genotype (N). 100 or more R7s were scored for each retina (n). Frequency was assessed using custom semi-automated software (see below) or manually. Error bars indicate standard deviation (SD).

### Image Processing

We employed a custom algorithm to identify the positions of individual R7 photoreceptors within an image of the fly retina. The script that implements our algorithm is available at https://app.assembla.com/spaces/roberts-lab-public/wiki/Fly Retina Analysis.

### Genome-Wide Association Studies

Genotype data from the DGRP freeze 2 lifted to the dm6/BDGP6 release of the *D. melanogaster* genome was obtained from (ftp://ftp.hgsc.bcm.edu/DGRP/). Phenotypes were calculated for the progeny of crosses of DGRP lines and *Df(3R)Exel6269* flies. To estimate genotypes of these flies from the DGRP data, we simulated each cross. For each SNP or indel variant in the DGRP genotype data, we assigned a new genotype: 1) homozygous reference remains homozygous reference, 2) homozygous alternate maps to homozygous alternate *if* in deficiency region, otherwise heterozygous, and 3) all other genotypes mapped to missing or unknown and not included in subsequent analyses. We performed quantitative trait association analysis using plink2 --linear (version 1.90 beta 25 Mar 2016; PMID:25722852). To reduce the impact of population structure, we included the first 20 principal components of the standardized genetic relationship matrix as covariates (calculated using plink2 --pca). To empirically correct p-values for each site, we performed a max(T) permutation test with 10,000 permutations (mperm option to plink2).

### CRISPR-mediated mutagenesis

*sin* was inserted into a lab stock line using CRISPR (Gratz et al. 2013; Port et al. 2014).

### T-maze Behavioral Assays

T-maze assays were conducted as described in (Yamaguchi et al. 2010).

### Consensus sequence

For the B1H data sets, WebLogo3 was used to generate position weight matrices (PWMs)(Zhu et al. 2011; Enuameh et al. 2013) (**Fig. 3B, S1A**). For the SELEX-SEQ data sets, MEME-ChIP version 4.11.2 was used to generate PWMs (Machanick and Bailey 2011; Nitta et al. 2015) (ENA: ERX606541-ERX606544).

### Conservation analysis

The Klu site and neighboring sequences for 21 *Drosophila* species were obtained from the UCSC genome browser. TOMTOM version 4.11.2. was used to generate the conservation PWM (Gupta et al. 2007) (**Fig. 3C, S1B**).

### SELEX-seq analysis

SELEX-seq datasets from (Nitta et al. 2015) were obtained from ENA (ERX606541-ERX606544). For read level analysis, we counted the number of reads containing the Klu binding site with *sin*, without *sin*, and neither site (there were no reads with both sites). We performed McNemar’s test to assess significance. We computed the frequency of each 10-mer within each dataset using Jellyfish version 2.2.6 (Marcais and Kingsford 2011). Using these counts, we determined the number of 10-mers with frequency greater than that of the Klu binding site with and without *sin.* Frequencies reported are for the combination of all four SELEX datasets.

### Population genetic analyses

We estimated allele frequencies from populations sampled world-wide at *sin* and at other 1-2bp indel polymorphisms. Allele frequency estimates based on pooled resequencing of populations sampled in North America and Europe were obtained from (Bergland et al. 2014) and (Kapun et al. 2016). Allele frequencies based on haplotypes (Lack et al. 2016) were also obtained from populations sampled North America, the Caribbean, Europe, and Africa.

